# Robust high-throughput phenotyping with deep segmentation enabled by a web-based annotator

**DOI:** 10.1101/2022.03.11.483823

**Authors:** Jialin Yuan, Damanpreet Kaur, Zheng Zhou, Michael Nagle, Nicholas George Kiddle, Nihar A. Doshi, Ali Behnoudfar, Ekaterina Peremyslova, Cathleen Ma, Steven H. Strauss, Li Fuxin

## Abstract

The abilities of plant biologists and breeders to characterize the genetic basis of physiological traits is limited by their abilities to obtain quantitative data representing precise details of trait variation, and particularly to collect this data at a high-throughput scale at low cost. Although deep learning methods have demonstrated unprecedented potential to automate plant phenotyping, these methods commonly rely on large training sets that can be time-consuming to generate. Intelligent algorithms have therefore been proposed to enhance the productivity of these annotations and reduce human efforts. We propose a high-throughput phenotyping system which features a Graphical User Interface (GUI) and a novel interactive segmentation algorithm: Semantic-Guided Interactive Object Segmentation (SGIOS). By providing a user-friendly interface and intelligent assistance with annotation, this system offers potential to streamline and accelerate the generation of training sets, reducing the effort required by the user. Our evaluation shows that our proposed SGIOS model requires fewer user inputs compared to the state-of-art models for interactive segmentation. As a case study in the use of the GUI applied for genetic discovery in plants, we present an example of results from a preliminary genome-wide association study (GWAS) of *in planta* regeneration in *Populus trichocarpa* (poplar). We further demonstrate that the inclusion of semantic prior map with SGIOS can accelerate the training process for future GWAS, using a sample of a dataset extracted from a poplar GWAS of *in vitro* regeneration. The capabilities of our phenotyping system surpass those of humans unassisted to rapidly and precisely phenotype our traits of interest. The scalability of this system enables large-scale phenomic screens that would otherwise be time-prohibitive, thereby providing increased power for GWAS, mutant screens, and other studies relying on large sample sizes to characterize the genetic basis of trait variation. Our user-friendly system can be used by researchers lacking a computational background, thus helping to democratize the use of deep segmentation as a tool for plant phenotyping.

## Introduction

Advances in high-throughput genome sequencing and computation have enabled the investigation of genomic variation within large populations [1–4]. In genome-wide association studies (GWAS), statistical models are used to model relationships between genotype and phenotype. When significant relationships appear, this can lead to the discovery of genetic markers and/or genes responsible for the phenotype of interest [3, 5]. However, genomic sequencing capacity and precision has outstripped the ability to produce high-quality phenomic data at scale. Phenotyping often requires individual inspection of large numbers of samples, with statistical power depending on sample size; for instance, as the sample size of a GWAS or mutant screen increases, the representation of rare genetic markers increases along with the power of models to discover them [6–8]. Sample size can be limited by the time cost of phenotyping, and both the accuracy and precision of measuring complex phenotypes as are common in studies of plant growth and development [9, 10]. The constraint of sample size therefore limits the insights that can be gained from these studies. Furthermore, phenotyping by humans commonly involves summarization of phenotypes in a manner that limits detail and subjects data to potential biases of individuals performing phenotyping. For example, samples may be given an ordinal or binary “score” representing a general summary of their complex trait or traits, which would be difficult to quantitatively measure in a simple and objective manner. For studies similar to our case example involving plant regeneration, common traits studied include callus size and quality [11], rates of callus differentiation into shoot [12] and the numbers of shoots produced [13]. These are very difficult to accurately measure without an annotation and imaging system, and their measurement will often compromise the required sterility of *in vitro* cultures during sequential analyses of growth. Comparable levels of complexity can be seen in populations of growing cancerous cells imaged with microscopy [14, 15].

While deep learning has demonstrated unprecedented abilities for high-throughput and precise phenotyping of complex plant traits including plant stress, disease, and development of specific tissues, these methods generally require large training sets that are labor-intensive to generate [16–19]. The amount of labor needed for phenotyping can be greatly reduced by supervised deep learning, although not eliminated altogether due to an inherent reliance on training data, which requires manual inspection and labeling of some number of samples.

Multiple approaches have been employed to alleviate the need for large training sets or to reduce the time cost of generating them. The number of training samples required for desired accuracy can be reduced, for example, by employing transfer learning. With this ubiquitous approach, users begin with a model trained for a similar task (often using a gold-standard training set) and retrain the model using a training set for their specific task. This re-training process tends to require relatively few annotated examples, whereas many more would be needed if the model were not already trained for a similar task [20]. Additionally, data augmentation approaches, such as those involving image rotation or other linear transformations, can be used to produce multiple training samples from each image [21]. Generation of training labels is particularly laborious when research designs require precise segmentation of image boundaries rather than simple classification of images. In a recent study, deep segmentation of healthy and cancerous tissues relied on a training set with precisely drawn lines separating healthy and cancerous tissue; the labor-intensive task of producing this training set was crowdsourced among 25 medical doctors and students [22].

As an alternative to manual drawing of segment boundaries, several established methods and tools for annotation depend on drawing of boxes or polygons [23–26], presenting a challenge for precisely labeling curved edges of objects such as leaves and other plant tissues. Our system features a user-friendly web-based annotator tool (IDEAS, Intelligent DEep Annotator for Segmentation) that applies deep segmentation to detect boundaries of objects given positive and negative cues by the user and a semantic prior map, thus enabling rapid and reliable labeling of objects with pixel-level precision (Fig. 1).

**Figure 1.**
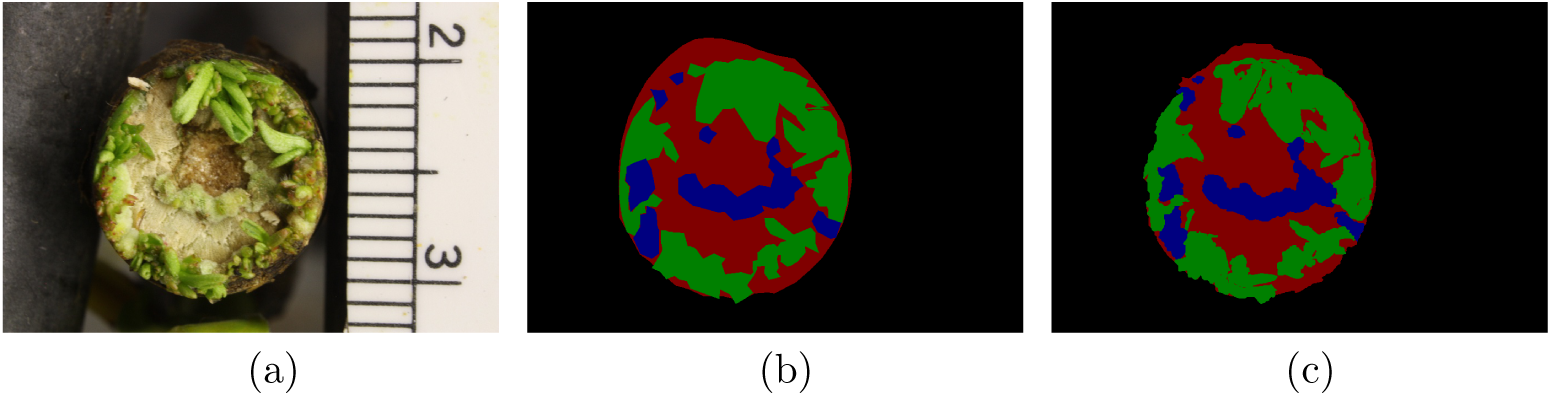
Annotation of an example image of a poplar stem tip undergoing regeneration: (a) Image to be annotated. (b) Polygon-based annotation from LabelMe, which was generated using 364 clicks precisely on the boundaries. (c) Annotation from IDEAS, which was generated based on 25 positive marks and 49 negative marks. This comparison demonstrates how IDEAS can be used to produce highly precise boundaries for objects with reduced effort.

Deep learning requires a massive amount of data to train a model. Generation of large datasets by human unassisted may prove time-prohibitive, especially for complex tasks. The time cost can be mitigated by the use of intelligent annotation algorithms to simplify image annotation tasks. Much research has been dedicated to building intelligent algorithms for automatic image annotation and building image annotation tools [27]. Nevertheless, these algorithms fail to predict well for complex structures and require a considerable amount of user effort to correct the incorrect model predictions. Hence, there is a need for an efficient tool for image segmentation using an intelligent algorithm that requires minimal user interaction and can label datasets belonging to any distribution of data. Our system, IDEAS, facilitates labeling highly precise curved objects with minimal labor and works well for segmenting previously unseen classes.

For the backend algorithm used in IDEAS, we propose to utilize semantic probability maps as a prior that acts as an additional guide to our interactive object segmentation model. The availability of strong prior information helps to guide the neural network in producing accurate boundaries and reduces the level of user interaction needed in providing positive and negative markers. Semantic segmentation has made tremendous progress in the last few years and is now among the fastest and most mature algorithms in deep learning. With state-of-the-art architectures such as DeepLab [28], PSPNet [29], and RefineNet [30] delivering strong performance on challenging benchmarks, obtaining semantic results for images is no longer difficult. Considering the potential for transfer learning to reduce the amount of training data needed, we hypothesize that the user effort required in providing clicks to guide segmentation can be reduced if segmentation is guided by a prior generated by a model trained on a partial dataset.

Leveraging semantic information to guide the main task has been previously followed in object detection [31, 32], dense object reconstruction [33] and interactive co-segmentation 34 tasks. However, to the best of our knowledge, our method is the first to apply this strategy for interactive object segmentation. We performed a benchmark experiment to evaluate the time cost advantage of using a semantic prior map along with the user click inputs, and found that use of the semantic prior map provides a substantial improvement over the use of user inputs alone.

The nature of our overall system as a web-based tool offers advantages for collaborative annotation and system administration. Multiple users can log into the system online from personal computers, working on a server with IDEAS installed and managed centrally by an administrator. Users benefit from the flexibility of being able to work from any computer with internet access, while the computationally intensive task of generating labels is assigned to a GPU on the server. Furthermore, users without a computational background benefit from an ability to apply the system without installing and updating software or otherwise interacting with the command line, as these tasks are handled centrally by an administrator.

After completion of a training set with IDEAS, a neural network is trained and used for semantic segmentation of large numbers of images that could not be practically annotated by humans. We use the state-of-the-art PSPNet architecture [29], notable for a global pyramid pooling module that enhances model accuracy. Finally, traits computed from segmented images provide quantitative measures of growth of distinct plant tissues. These measures include the relative areas and counts of specific tissues, thus providing an array of statistics biologists may choose from to represent tissue development in downstream statistical models.

## Materials and methods

IDEAS (Fig. 2) features separate front-end and back-end components. The front-end, accessible to the user through a web browser, contains all user-facing, interactive modules. These modules cooperate with each other to support the workflow and are implemented through HTML, JavaScript, and jQuery. On the back-end, the Python Flask web framework is used to receive requests from the front-end and return the generated object mask for the object being annotated. *IDEAS is deployed at https://ideas.eecs.oregonstate.edu and is accessible to the public*.

**Figure 2.**
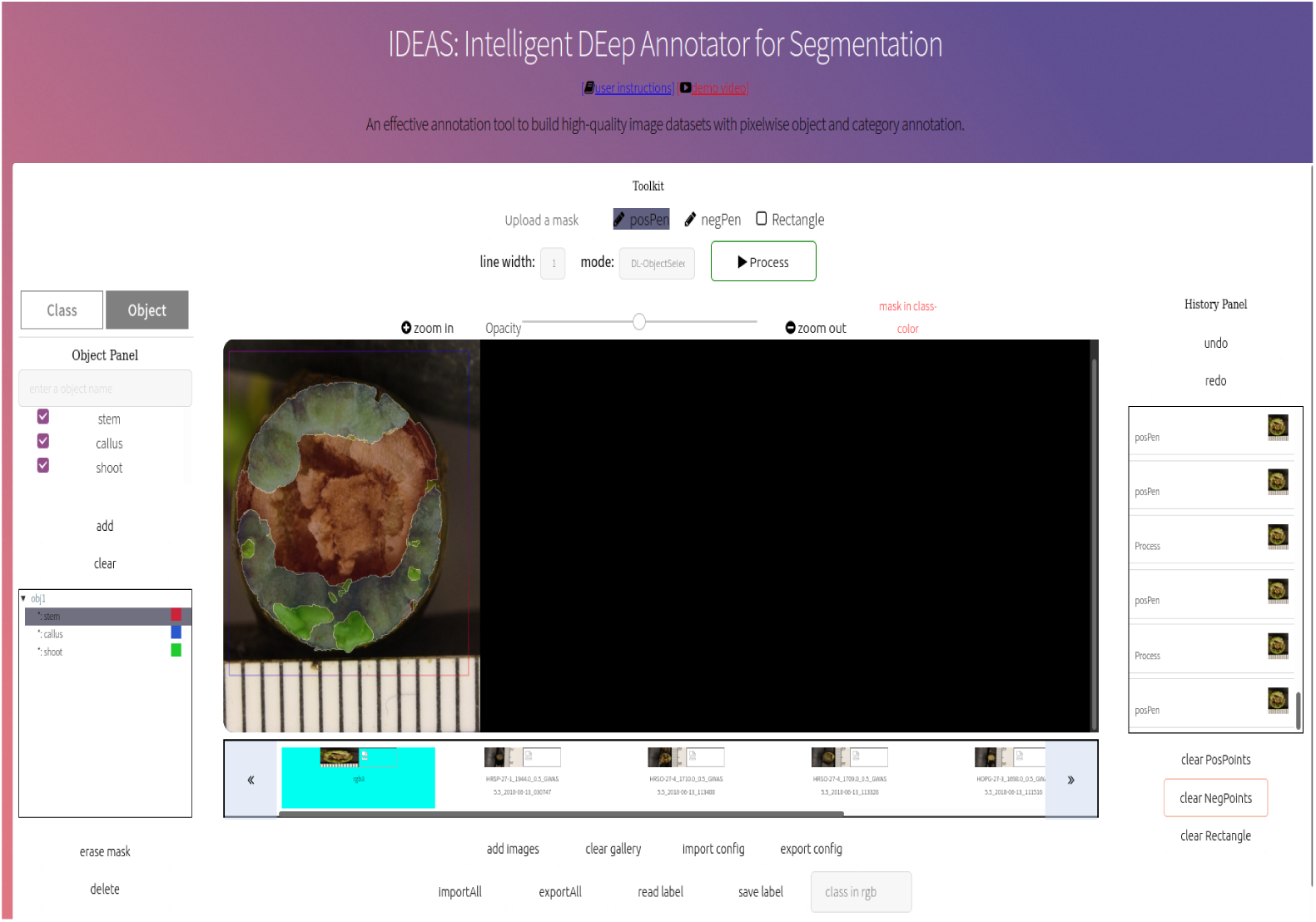
Screenshot of IDEAS with one annotated image in a web browser

The development of IDEAS enables the collection of ground truth segmentation masks, to be used as training labels, for various types of plant images. A semantic segmentation model can then be trained using the collected annotations to compute masks for large sets of unseen images. Below, we introduce features of front-end, back-end, and auxiliary functions on IDEAS.

### IDEAS: Front-end Interactive Modules

The front-end interface of IDEAS features seven different modules(Fig 3):

**Figure 3.**
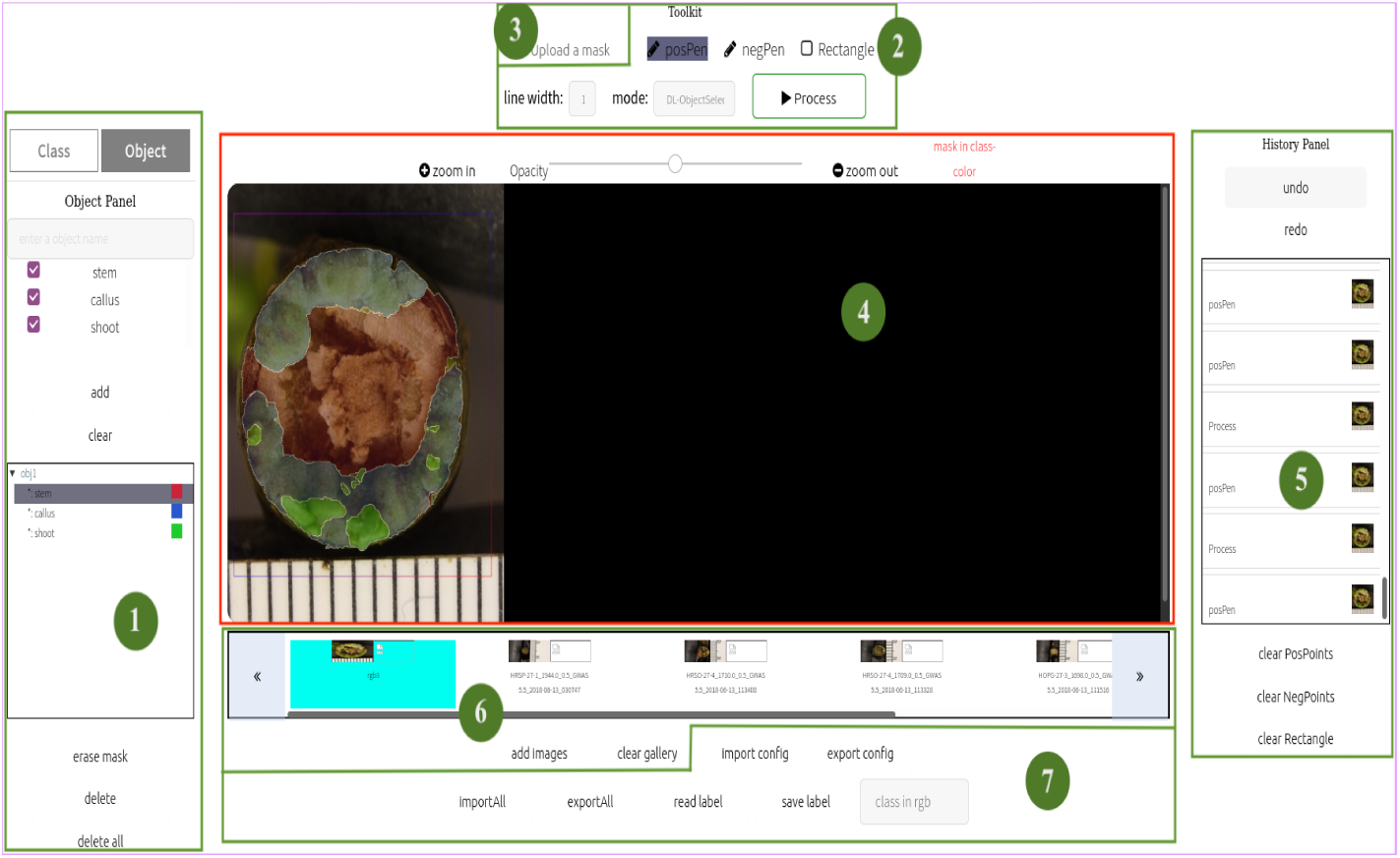
Front-end modules in IDEAS

### ➀ Object and Class Panels

Our system allows the user to customize the identity of objects and the classes they include. Fig 4(a) shows the unfolded class panel and object panel. The class panel displays a list of classes to be annotated. Using the object panel, the user can define objects and their nested classes by either 1) adding multiple classes to an object when defining the object on the object panel, or 2) using the button ‘add to’ on the class panel to add classes to a specified object one-by-one. The intuitive design of object and class hierarchy is influenced by applications that involve analysis of multiple, distinct parts of given objects (e.g. distinct plant tissues such as callus or shoot, growing in each individual sample on a petri dish (Fig.4(b))).

**Figure 4.**
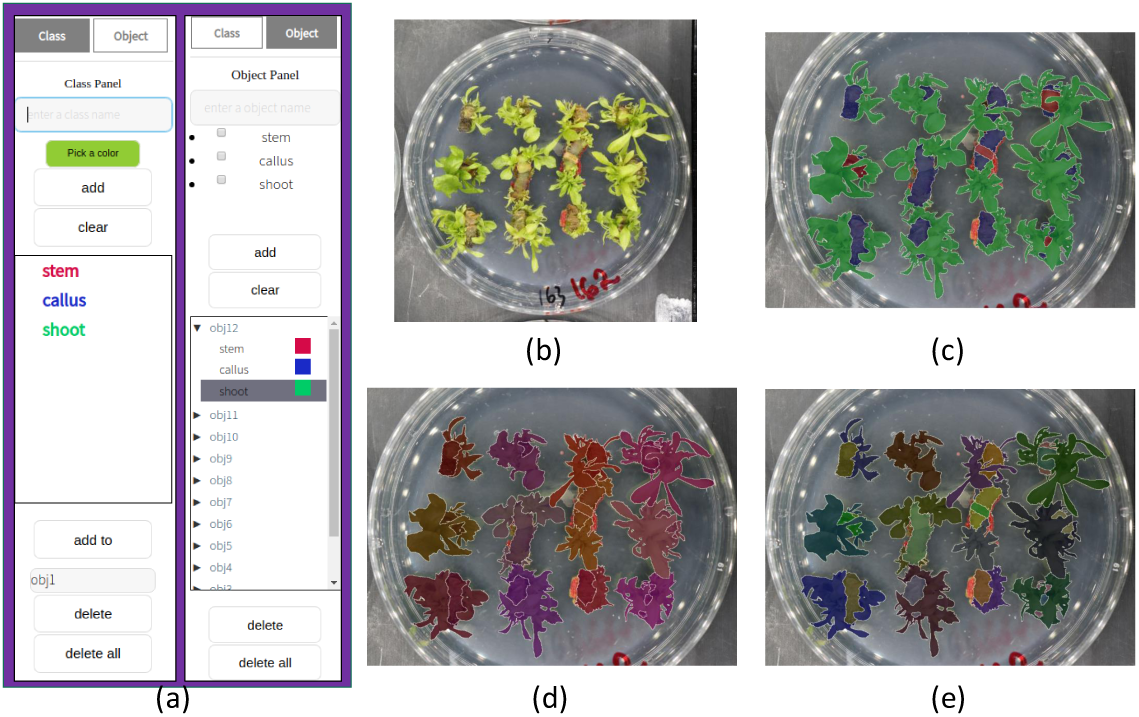
One annotation example: (a) Customized configuration of class and object hierarchy. (b) RGB image. (c) (e) RGB images with superimposed labels for class (c), object (d), and nested object-class (e).

### ➁ Toolkit

Point-and-click tools for user annotation of images include: 1) Two types of pens, for drawing positive and negative strokes inside and outside of a desired segment, respectively, which are used by the back-end to generate the object mask; 2) A bounding box tool to specify a local window around a specific object the user desires to interact with, accelerating computation time by limiting mask computation to pixels within the box. An alert is generated recommending the use of this tool when annotating images larger than 512 × 512; and 3) drop-down menus allowing the user to select line width for marks drawn and to choose between manual annotation and a deep algorithm for intelligent boundary detection.

### ➂ Mask Upload

This tool allows the user to upload a semantic mask to provide prior information to the classifier, which can reduce the number of user clicks required for strong performance in our deep interactive model, SGIOS. If no mask is uploaded, the algorithm utilizes a blank mask and produces results equivalent to those from interactive segmentation without a semantic prior map.

### ➃ Canvas

The central editor, implemented as an HTML5 canvas element, provides a space for the user to apply tools from the toolkit (rectangle and positive/negative strokes) on a given image. Zooming functions can be used to assist in fine-scale drawing or to provide a broad overview of an image. The user begins interaction with the canvas first by selecting a nested class for an object of interest (from the object panel), then draws positive and negative strokes and clicks the ‘process’ button in the toolkit module. For quick and efficient labeling, these strokes are usually drawn as short lines. Each pixel in a stroke is considered an example of a pixel inside (positive strokes) or outside (negative strokes) the segment of interest. The user-provided marks are sent to the back-end, which detects boundaries and returns a mask to be superimposed over the segment of interest in the graphical front-end. This superimposed mask appears with a white line on boundaries, with inner pixels visualized with a user-selected level of opacity (set using the opacity ranger), allowing the user to see both masks and underlying objects. If the mask does not provide an adequate fit for the desired segment, the user can provide additional positive and negative strokes and again click the ‘process’ button to generate a new mask.

### ➄ History Panel

A history panel is integrated into the annotator to help the user recognize and correct errors. Users can view independent steps of annotation in this panel and correct mistakes with undo and redo functions. The history panel also provides a ‘clear’ button to erase all operations and return to the initial state. Three additional buttons allow the user to undo specific toolkit actions. These include ‘clearPositivePoints’ and ‘clearNegativePoints’ buttons to erase all ‘posPen’ and ‘negPen’ marks, as well as a ‘clearRectangle’ button to remove the bounding box set with the rectangle tool.

### ➅ File Gallery

A file gallery is displayed underneath the canvas, providing an overview of all uploaded images. Images can be uploaded by two methods: first, through the image upload prompt seen when logging into IDEAS, or alternatively, using the ‘addImage’ button to add images as annotation is ongoing. Using either method, images can be uploaded either independently or in batches and their thumbnails are displayed in the file gallery. To switch the image being annotated in the ‘Canvas’ panel, the user can simply click the thumbnail of another image. To remove all uploaded images from the gallery, the user clicks the ‘clear gallery’ button.

### ➆ Importing and Exporting

Upon completing annotation for a given image, the user can save the labels (generated mask) as a PNG image file with labels recorded in any of three formats: with segments colored by either their class or object, or both. An example is shown in Fig.4(c)-(e). The customized class and object hierarchy information can be exported as a .txt file and re-imported during later annotation sessions, a quick and simple means of maintaining consistent settings over the course of an experiment. Furthermore, the user has an option to import or export a .txt file containing the entire annotation environment, including all user actions and generated masks. This feature enables the user to restore the environment to update annotations at a later time.

### IDEAS: Back-end deep interactive algorithm

When the user clicks the ‘process’ button, the front-end collects user inputs including positive and negative strokes, the prior mask, and the bounding box, along with the image and any pre-existing labels, and sends this data to the back-end. The back-end then generates a mask using one of two algorithms, as selected via the ‘mode’ drop-down menu under toolkit. In *manual* mode, the pointer is used to mark every pixel of the desired segment. Annotating whole images by this method can be difficult and time-consuming, as the user is required to draw a label covering the segment of interest and has no assistance in accurately drawing boundaries. Contrarily, the *DL-ObjectSelect* mode which uses SGIOS provides a rapid means of automatically segmenting the selected object when the user provides a small number of positive and negative strokes. We demonstrate that this latter method is fast and efficient, minimizing the required inputs and time commitment from the user. Thus, the *manual* mode is rarely used in practice.

### SGIOS (Semantic-guided Interactive Object Segmentation)

Segmentation annotation of images presents a challenge in that every pixel must be labeled for each training image, which could be a time-consuming process, especially in plants where the boundaries usually cannot be expressed with simple geometric shapes. Intelligent algorithms are needed to reduce the human effort required to produce these labels. Here, we propose an interactive segmentation algorithm utilizing a semantic probability map as prior information to guide the segmentation while the user provides inputs of positive and negative clicks. The use of prior information is a strategy to guide the deep convolutional network and provide context for the object of interest, reducing the number of user inputs required for desired performance.

Fig. 5 illustrates the pipeline for computing object masks after an object of interest is labeled. The image, user-interaction pairs (positive and negative marks), and semantic information are used as inputs for a fully convolutional network (FCN), which is trained with the bootstrapped cross-entropy loss function to predict a binary mask of the object. On the user interaction guided map, we calculate the pixel value *u*_*x,y*_ at location (*x, y*)as the minimum Euclidean distance between (*x, y*) and the set of marks. In other words, given a set of points *P* = {(*i, j*)}, where (*i, j*) is the point location, *u*_*x,y*_ = min_(*i,j*)∈*P*_ (*i* −*x*)^2^ + (*j* −*y*)^2^

**Figure 5.**
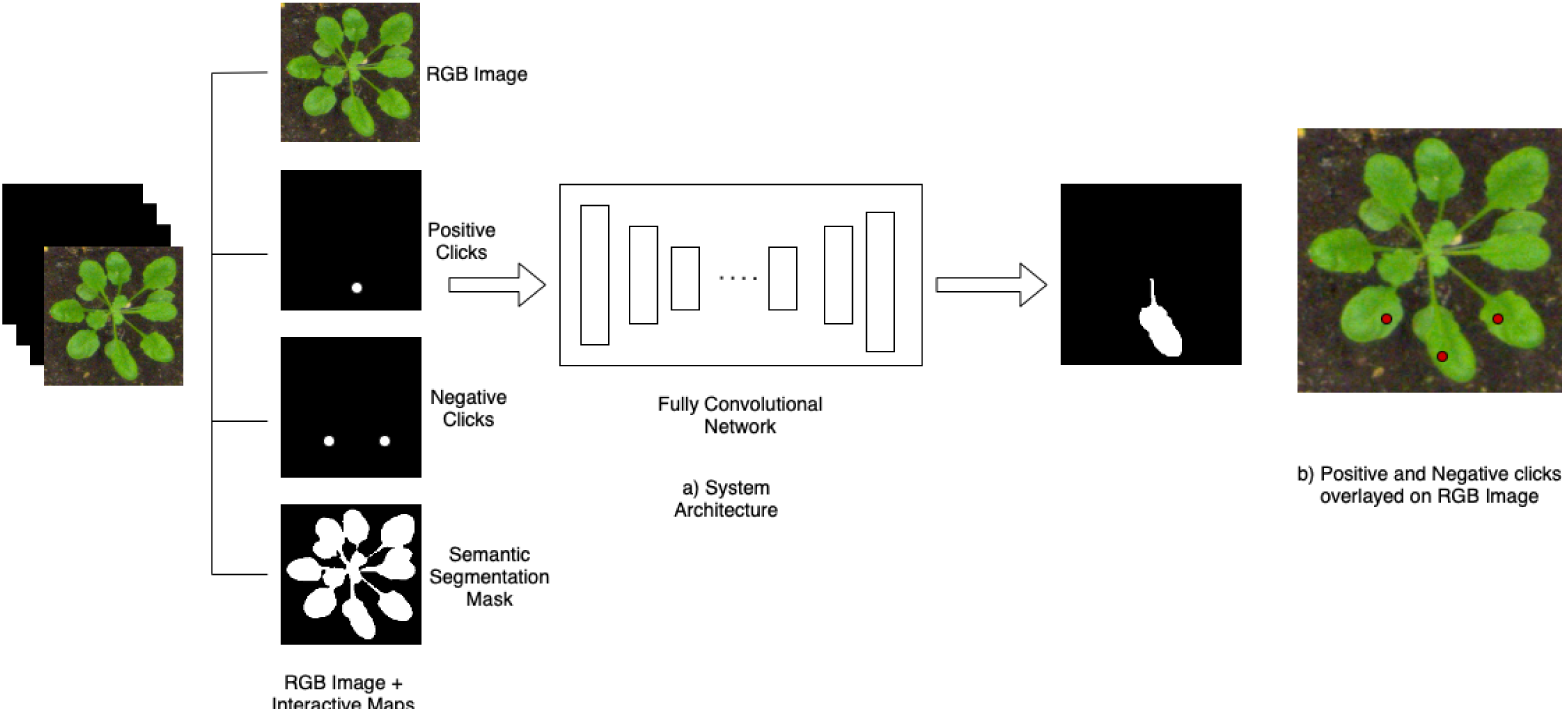
Pipeline of the Semantic-Guided Interactive Object Segmentation Algorithm

Most CNN networks [35] for interactive object selection are trained non-iteratively, despite the often iterative nature of refining results with additional clicks by the user. Following the iterative training strategy proposed by ITIS [36], we choose to train our model iteratively. Simulated initial user clicks are updated with the corrective clicks based on the mask obtained in the previous iteration, and this information is sent to the FCN as an input to predict the refined mask. Interactive clicks are further simulated until the desired mask is finally obtained. Below we describe the key components in training our interactive model, SGIOS, with simulated user clicks.

- **Initial click sampling:** To make our model flexible and not depend on the location or the number of clicks, we randomly sample *n* positive and *m* negative clicks until a total of 20 clicks are placed. To sample the positive clicks, we begin by identifying points in the center of the object and randomly sample a point from these. All positive sampled points lie on the object of interest and are d_step_ away from the previously sampled positive points. We first sample negative clicks on other objects close to or touching our object of interest, and the remaining clicks are sampled around the object. When the number of negative clicks sampled is zero, we send a blank mask for the negative clicks and a positive click encoded mask with clicks only on the object of interest.
- **Iterative click sampling on incorrect prediction:** Our iterative click sampling strategy is similar to that proposed by ITIS [36], with modifications in the erroneous region selection and click placement on each iteration. Below we describe the steps for generating the correction clicks while training.
  - The output prediction from the last step is compared with the ground truth mask for the object of interest, and the incorrect pixels are grouped into multiple clusters using the connected components.
  - Instead of selecting the largest cluster on each iteration as is done with ITIS [36], we sample R regions where the clicks should be placed from the distribution of all regions. The distribution of regions is weighted by the probability proportional to the number of pixels in each region.
  - K points are randomly sampled from the R selected erroneous regions, where K*>*=R and the total number of clicks is thresholded to 20.
  - The selected clicks are encoded similarly as the initial clicks, using a Gaussian distribution. An example of iterative click sampling is shown in Fig. 6.

**Figure 6.**
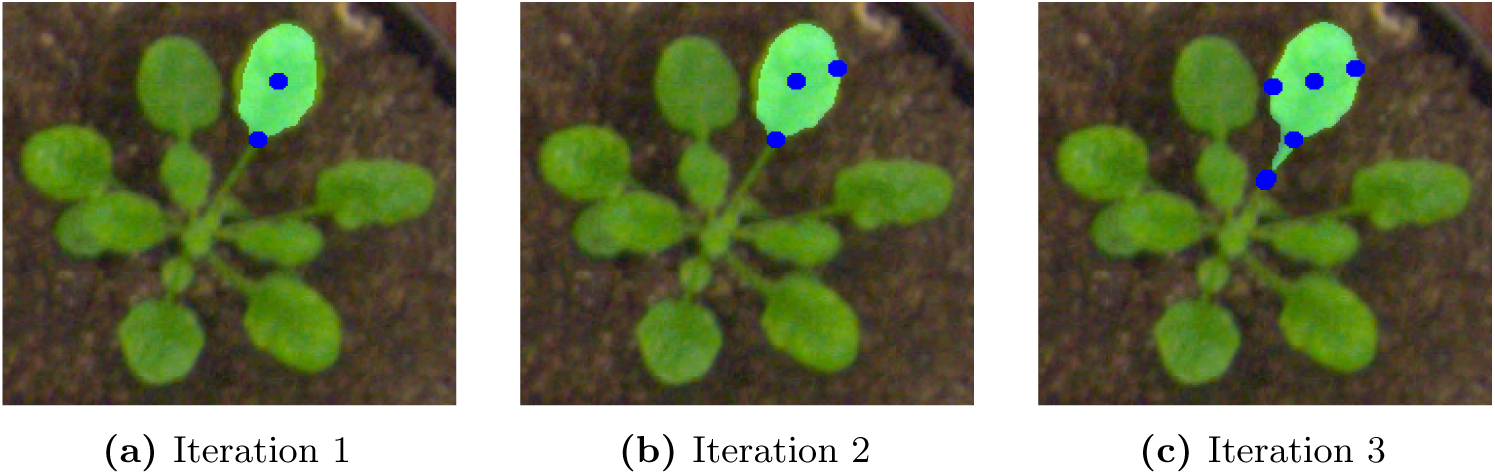
Iterative click sampling on incorrect predictions. (a) Initial prediction when two clicks are placed on the object of interest (b) Prediction after an additional click is sampled in the right error region (c) Prediction after one click is sampled in each of the remaining two error regions
- **Semantic Prior Maps:** The semantic prior maps were generated from an offline semantic segmentation model. For example, in our case study, we used a Deeplabv3+ model trained on plant images for segmenting a image into three categories (stem, callus, and shoot) along with background. The multi-channel output before the final sigmoid layer in the semantic segmentation model was output as the semantic probability maps featuring one channel for each class. Since the semantic prior maps are floating-point numbers on each pixel, we saved them in TIFF format. To be noted, the users can use a pre-trained semantic model that was trained for the categories of interest to generate these prior maps. IDEAS does not support online semantic model training or generating TIFF files. Users should handle this outside of the system.
  - The appropriate semantic channel is selected based on the semantic category or categories of positive clicks. With the assumption that the user will always place at least one positive click on the object of interest, we obtain the semantic probability map corresponding to the category with maximal probability at the given pixel locations. Using this approach, we can adapt to semantic segmentation tasks with any number of categories. With inclusion of the semantic prior map, we are often able to obtain adequate results for interactive object segmentation with only a single positive click on the object.
  - When no semantic prior map is provided, interactive object segmentation proceeds without this information, using a blank mask. This approach is most appropriate when producing a training set for an initial model, which can be used for producing the semantic prior maps for the remaining dataset.

### IDEAS: Auxiliary functions

Beyond the fundamental workflow introduced above, we emphasize the following features in IDEAS to enhance precision, usability and annotation speed:

- **Positive/Negative Strokes** We incorporated a feature to allow the user to provide groups of positive or negative markers as strokes as an alternative to providing single markers one click at a time. Rather than only clicking to provide individual points as positive and negative marks, the user can move the mouse while holding the click, drawing strokes to mark multiple pixels along a path. The ability of the user to quickly provide many marks is valuable because segmentation accuracy depends on the number of marks drawn.
- **Quality improvement by adding more ‘posPen’ or ‘negPen’ strokes**: We enhanced the semantic-guided interactive object segmentation by adding a feature that enables the user to improve the returned object mask by adding additional positive and negative strokes and again pressing the ‘process’ button. This ”refinement” process is integrated into IDEAS. As shown in Fig.7, the accuracy of object mask boundaries improves as additional positive/negative strokes are given by the user. This provides an easy and robust means for the user to produce annotations with desired boundaries precisely labeled.

**Figure 7.**
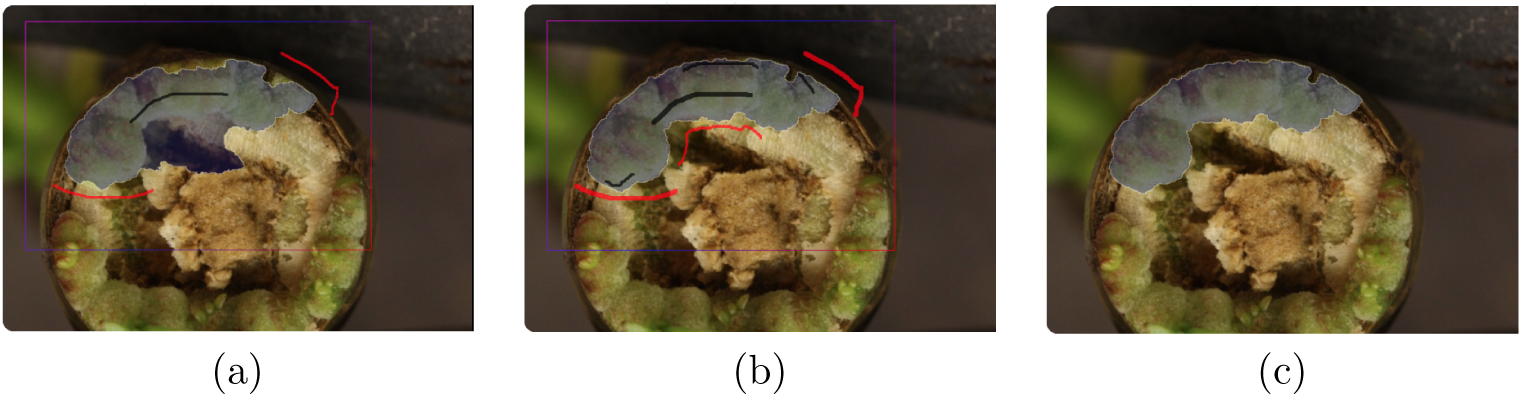
Demo of the ”refinement” process: (a) The labels from given ‘posPen’ and ‘negPen’ strokes is errant along the boundary. (b) User adds more strokes and the precision of the predicted labels is highly improved. (c) Finalized label for the object of interest.
- **Lock/Unlock the labeled objects**: The user can lock or unlock each single labeled nested class for an object by double-clicking it on the object panel. As Fig.8 shows, the labeled objects cannot be modified while locked. Unlocking is necessary before the given object can be edited again. This locking function allows the user to annotate objects without affecting previously drawn labels of other objects. This is particularly useful when small objects are annotated prior to larger, nearby objects. The former objects can be locked as additional objects are labeled, then unlocked and edited again at a later time.

**Figure 8.**
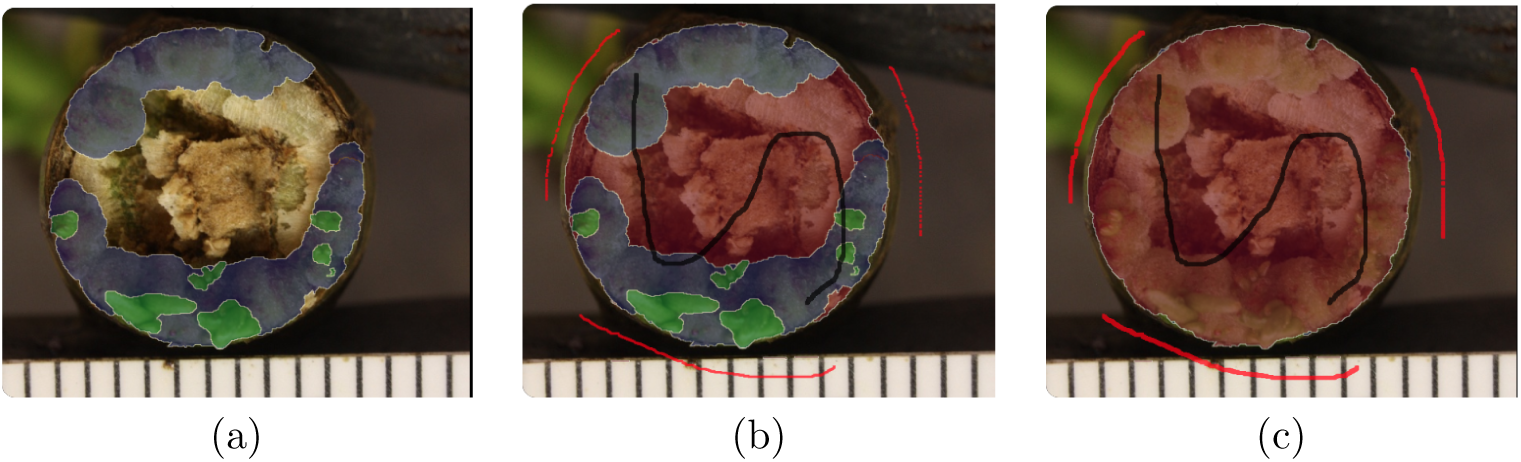
Demo of lock/unlock functions. (a) The classes *callus* (blue) and *shoot* (green) have been already labeled. (b) If *callus* and *shoot* are locked, annotation of the new object, *stem* (red) cannot change their labels. (c) If *callus* and *shoot* are unlocked, annotation of *stem* changes their labels.

## Results

In this section, we first showed the efficiency and accuracy of SGIOS on the Pascal VOC [38] dataset and the Computer Vision Problems in Plants Phenotyping (CVPPP) leaf segmentation challenge (LSC) dataset [39, 40]. Then we showed the performance of IDEAS with conducting a preliminary study with the help of an expert biologist, who helped perform annotations using our GUI. Finally, we presented an application of IDEAS for high-throughput phenotyping as part of a GWAS to identify genes controlling regeneration in poplar.

### Semantic-guided Interactive Object Segmentation

#### Implementation detail

We used the model architecture proposed by ITIS [36] with the modification of adding a semantic probability map input. To benchmark this approach, we simulated the user clicks in training using the previously described strategies for initial click sampling and iterative click sampling on incorrect predictions.

#### Training SGIOS on the augmented Pascal VOC dataset

We used the augmented Pascal VOC [38] dataset with additional annotation from the Semantic Boundaries Dataset (SBD) [41], which consists a total of 10,582 images for around 25,000 objects belonging to 20 different classes in the training subset. The validation subset consists of 3,500 object instances and was used for testing. We finetuned the SGIOS model for 50 epochs with a training batch size of 5 with an initial learning rate of 1*e* −4. The learning rate varied between the epochs using an exponential decay schedule with a decay rate of 0.9 every 5 epochs and was capped to 5*e* −7. To make the model flexible and avoid over-fitting to the semantic prior map, we randomly reset this prior map to a blank mask containing all zeros in the training stage.

#### Comparison with the state of the art models

We used the same methodologies used in prior work [36, 42] to evaluate our model’s effectiveness on Pascal VOC.

1. *The average number of clicks required to reach the IoU threshold* 85% *on Pascal VOC*. For each object, we compute the number of clicks used until obtaining a segmentation with intersection-over-union (IoU) larger or equal to 85%. If the desired IoU was not reached for any object within 20 clicks, we thresholded the number of clicks to 20. The average number of clicks was computed over all objects in the validation dataset. Table 1a reports the result, SGIOS achieves the best number of clicks at 3.1.

**Table 1.**
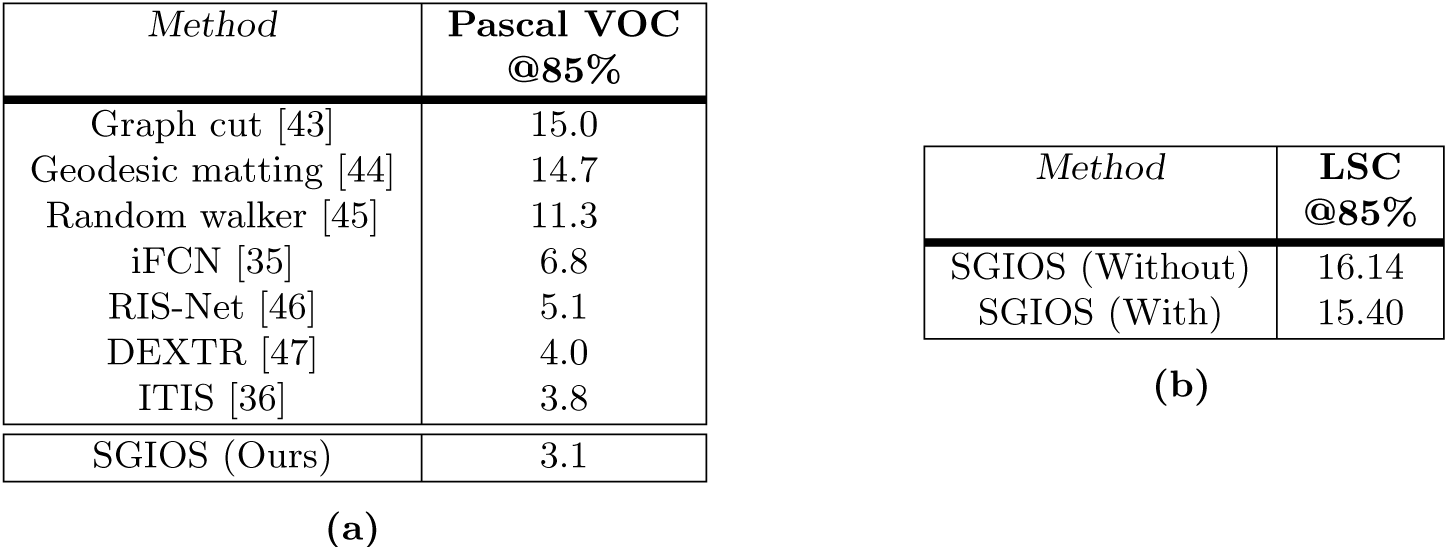
(a) Comparison of interactive models to find the number of clicks required to reach a threshold IoU (85%) on Pascal VOC. (b) Number of clicks required by SGIOS to reach a threshold IoU (85%) on the LSC dataset with and without using the semantic prior map.
2. *The mIoU with different numbers of clicks*. Fig. 9 shows the mIoU using different numbers of clicks over the objects in the validation dataset. This shows the benefit of using a semantic prior map is most apparent when a smaller number of clicks is provided. With a single click, SGIOS reaches an IoU of 74.7% and outperforms all other models by a considerable margin, demonstrating that the use of SGIOS reduces the number of clicks required for strong performance, thus mitigating the time cost of annotation. Figures S1 to S3 show several example predictions obtained on Pascal VOC using a single click, two clicks, and three clicks.

**Figure 9.**
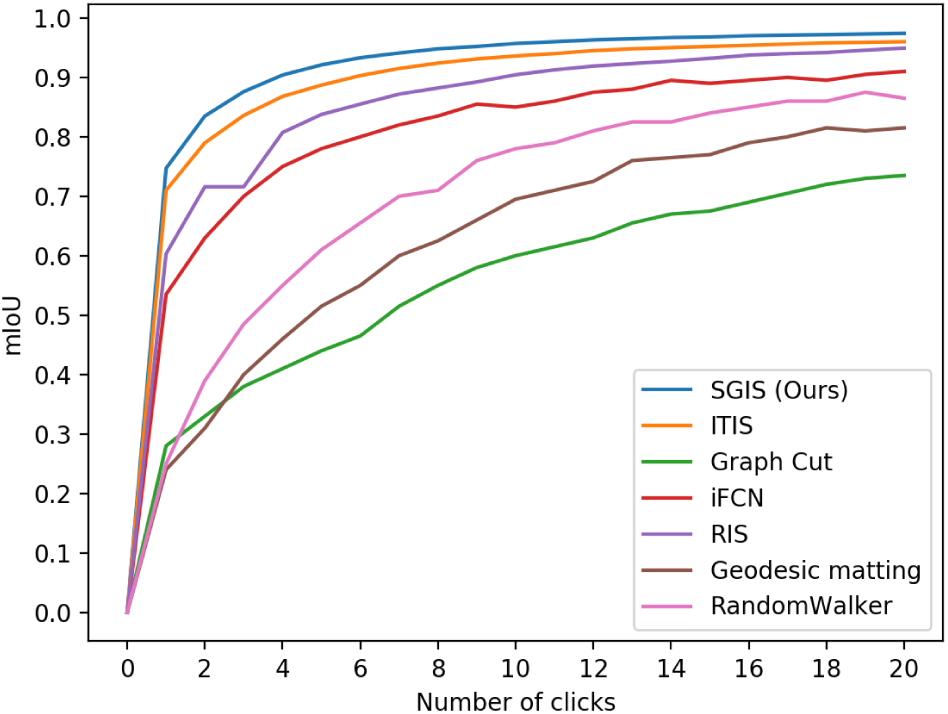
Mean IoU vs Number of clicks on Pascal VOC (Thresholded to 20)

#### Testing on the LSC dataset

We tested our SGIOS model on plant segmentation using the LSC dataset [39, 40]. We used 276 LSC images with object mask labels out of the 347 images provided in the original dataset. We considered every leaf to be a single object and obtain 3152 object instances from the 276 images. Table 1b demonstrates that using the semantic prior map provides a marginal benefit. The LSC dataset presents a particular challenge for instance segmentation because the leaves are in very close proximity and often obscure one another. Figures S4 and S5 shows some example predictions obtained on this dataset using two and three clicks, respectively. Please note that we used the model trained on Pascal VOC to test on this dataset.

#### IoU when semantic information is not present

We tested our model’s performance in the absence of semantic prior map by passing a blank mask to the semantic mask channel input. The average number of clicks required to reach the threshold of 85% for this subset is 3.5. This experiment confirms that the model is flexible to the semantic information and can perform reasonably well even if the semantic information is not present.

### A study on annotation using IDEAS

We conducted a preliminary study with the help of an expert biologist who helped perform annotations using our GUI. The study was performed to evaluate the advantage of using the semantic prior map for interactive image segmentation task. In particular, the study was designed to determine whether the use of a semantic prior map helps to save time to annotate an image.

#### Design of the study

A set of ten *in vitro* poplar tissue culture images were selected for annotation using IDEAS. These images were randomly divided into two counterbalanced groups as follows:

- Group 1 - The operator annotates five images without using the semantic prior map first and then annotates the same set of images using the semantic prior map.
- Group 2 - The annotator annotates five images using the semantic prior map first and then annotates the same set of images without using the semantic prior map.

The study took place in four sessions, with two sessions for each group. Each session took close to two hours to complete and was conducted over two days. The counterbalancing scheme was designed to control for the risk of bias from the learning effect when the operator annotates a given image twice and may recall details from the first annotation. Annotations were performed by an expert in plant issue culture. The participant was briefly introduced to how to use the system and was given time to practice using the system for a day before beginning the study. Time taken to annotate each individual explant on each plate was recorded.

#### Collected images

The images used were sourced from an ongoing study of *in vitro* regeneration in poplar (Fig. 4), a departure from our use of *in planta* samples presented for the earlier GWAS demonstration (Fig. 1, 2, 3). The complete dataset features a total of 1,278 genotypes, each represented by four plates with twelve stem explants. Two plates of each were subjected to an indirect regeneration treatment (callus induction media followed by shoot induction media) and a direct regeneration treatment (shoot induction with no pre-incubation on callus induction media).

#### Prepare the offline model for compute semantic prior map

We first randomly sampled 100 images from the collected dataset and annotated them on IDEAS with no semantic prior map, in which the default SGIOS model trained on Pascal VOC is used. We then fine-tuned the model Deeplabv3+ [28] for semantic segmentation. We use the data augmentation strategies (including random rotation, random flipping, random cropping) to bring a performance boost. The model was fine-tuned for 25,000 iterations using an SGD optimizer with a learning rate of 1*e* − 3 and batch size of 4. Afterwards, this trained model was used to generate semantic prior maps to assist further annotation using SGIOS, thus enabling accelerated expansion of the training set beyond the initial 100 images.

#### Timing of annotation

We recorded the time taken to annotate individual explants on the plate with and without semantic prior maps. The study consisted of annotating ten plates where each plate had 12 explants, yielding a total of 120 paired data points. We next performed statistical analysis to inform whether the difference in annotation speed results from an advantage of SGIOS, or is due to random chance. We converted the recorded time into seconds and checked the difference in time for normality using the Shapiro-Wilk test. By observing the histogram of the difference in Fig. 10 and the p-value from the Shapiro test equal to 1.868*e* − 12, we concluded that the data violates the normality assumption. Thus, we conducted a non-parametric test, the Wilcoxon signed-rank test, on the paired data. The Wilcoxon signed-rank test indicated that significantly less time was needed for annotation when semantic information was used, (V = 1954.5, p-value = 9.168*e* − 06, Wilcoxon signed-rank test). Boxplots for both groups are shown in Fig. 11. The mean time taken to annotate an explant using a semantic map is 19.99% (27.66 +/- 8.15 secs, n=120) less than the time taken to annotate an explant without using it.

**Figure 10.**
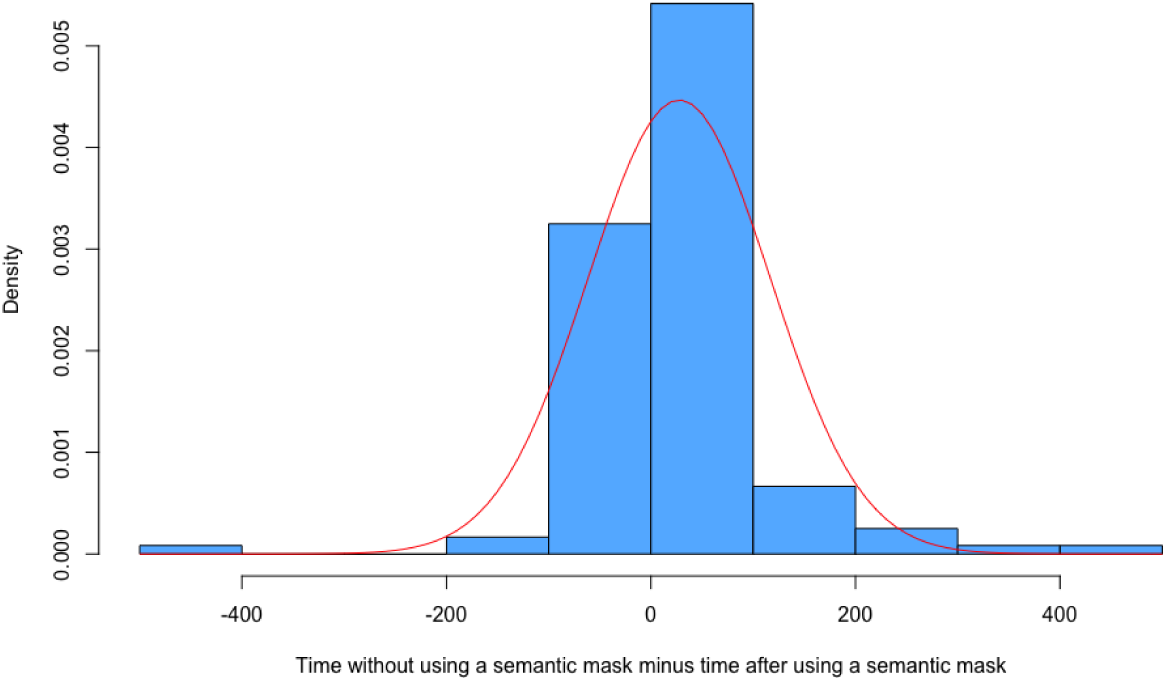
Histogram of difference in time with and without using the semantic mask, with time shown in seconds

**Figure 11.**
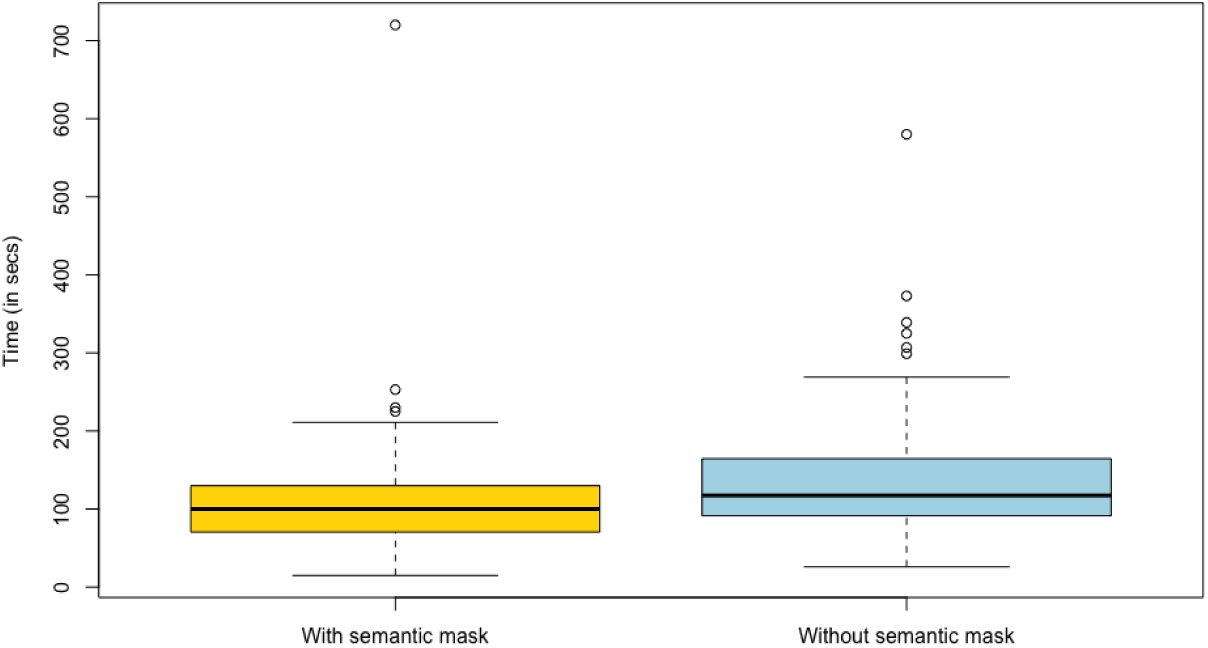
Time taken to annotate with and without the semantic mask (in seconds)

### A high-throughput phenotyping system with IDEAS

We applied our system, IDEAS, as part of a GWAS of *in planta* regeneration in *Populus trichocarpa* (poplar). In this experiment, we applied our phenotyping system toward discovering genes that contribute to variable regeneration response. Plant materials phenotyped included 1206 clones that represent genetic diversity from California to British Columbia and have been propagated at a field trial location in Corvallis, OR. Plant cuttings from a wide variety of natural genotypes [48] underwent a regeneration-promoting hormone treatment at the cut surface while rooting in water, then were imaged to provide RGB data for analysis. A subset of these images was annotated using IDEAS to build a training set. A semantic segmentation model was trained and then used to segment the remaining, unseen images. Using image processing techniques, we calculated statistics summarizing the segmentation outputs. These statistics were used to represent phenotypes in GWAS, revealing genetic markers associated with these phenotypes.

#### Phenotype data collection and summarization

- *Imaging*. The stem cuttings were collected from the field and incubated at 4° C for 2-4 weeks. Dormant stem cuttings were placed in 50mL Falcon tubes with water for five weeks. Once-per-week from the second week onward, the cut tip of each stem was treated with a droplet containing the plant growth regulator thidiazuron at 0.5mg/mL in water. Stem tips were imaged at weekly timepoints by conventional photography using a consumer-grade camera held over plants by a mount. As phenotyping all cuttings at once would have been impractical with limited resources, cuttings were divided into seven groups of up to 400 cuttings each, with groups undergoing this process one after another. This process was performed for up to 400 cuttings at a time. The first group was the only one to include images at the first week and did not include images for fourth and fifth weeks. The third group was missing data for the fourth week. From the fourth group onward, plants were studied in replicates of two and the computed values used for GWAS are the average of values for each replicate.
- *Annotating*. We collected 4896 phenotype images as described, then randomly selected 249 images and annotated them using IDEAS. Fig. 12 shows some examples of the annotated labels. Images were annotated to divide plant tissue into segments yet to undergo regeneration (stem) or one of two stages of regenerated tissue (callus or shoot). These annotated images were then randomly split into the validation set and the training set with 24 and 225 images, respectively. In Table 2, we summarized the content of the labeled dataset.

**Table 2.** Distribution of annotated images.

| Items | analysis result |
| --- | --- |
| Distribution over weeks 1~5 | 12.8%, 19.0%, 27.7%, 16.9%, 23.6% |
| Avg. percentage of annotated pixels | Shoot: 0.14%, Callus: 2.59%, Stem: 1.94% |
| Avg. number of isolated clusters | Shoot: 0.14, Callus: 3.00, Stem: 3.72 |

**Figure 12.**
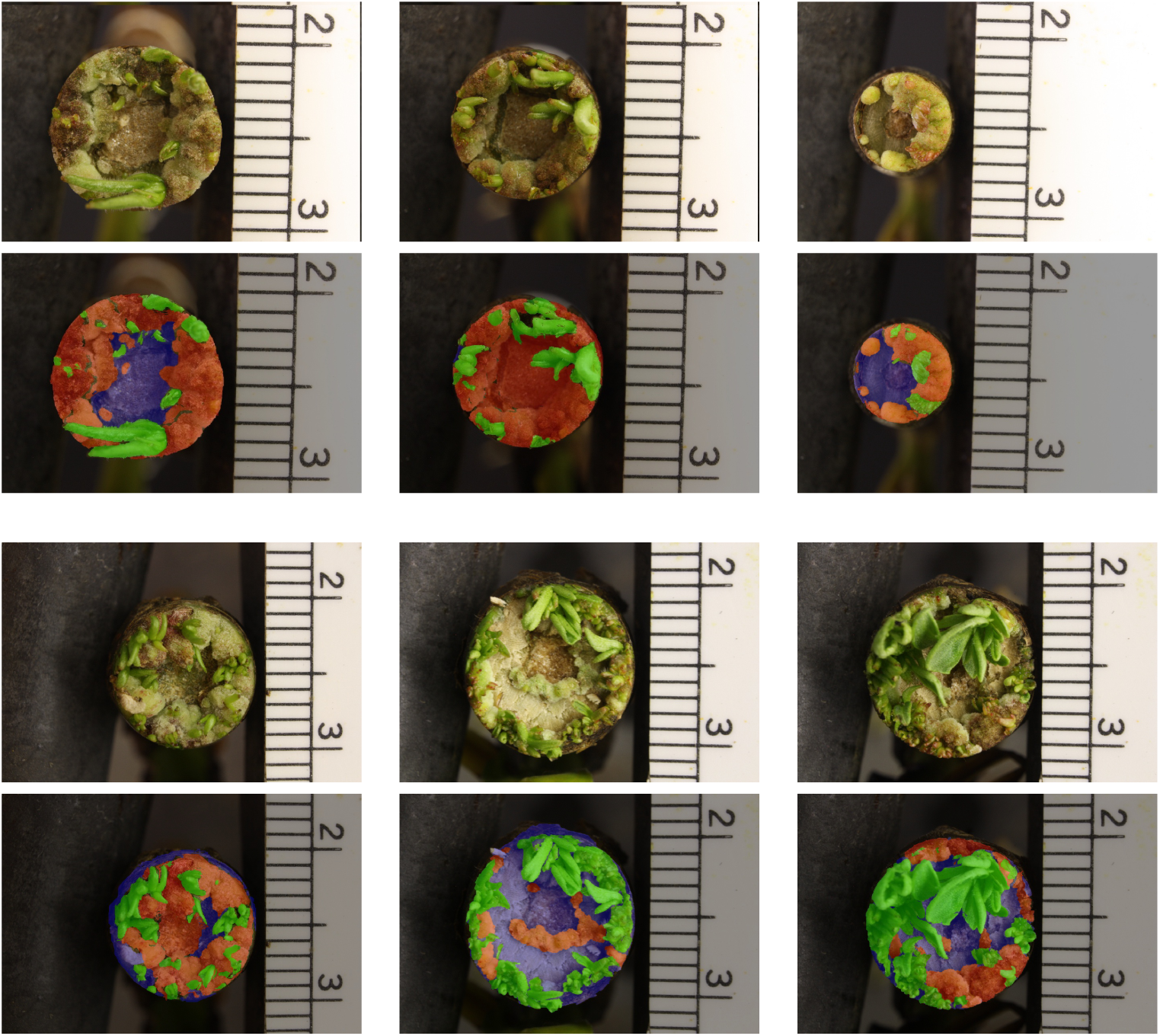
Examples of annotated ground truth labels overlapped with the RGB image: callus is in red, shoot is in green, and stem is in blue.

#### Plant Segmentation

We then trained a semantic segmentation model with the annotated images for computing biological traits. The details of this step are below.

- *PSPNet [29]*. PSPNet is among the state-of-the-art architectures used for semantic segmentation. We adopted PSPNet as the deep segmentation model and used *ResNet-50* as its backbone network in our work and employed the following training protocol to train the model to segment the images: We set the initial learning rate to 0.002 and decreased it using a polynomial learning rate schedule with power at 0.9 in each epoch. The training set was augmented with 1) random scaling to different sizes with scales 0.55 ∼ 1.0, 2) flipping horizontally with probability at 0.5, (3) random rotation with angle between −10° ∼ 10°. We trained the model for 60 epochs with batch size of 4 and weight decay of 4*e* − 4. To be noted, PSPNet can be replaced with other semantic segmentation model without affecting the phenotyping system pipeline.
- *Metric*. To verify the performance on semantic segmentation, we apply the IoU (intersection over union) metric, which is computed as 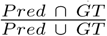. Here, we report the IoU for each class as well as the mean IoU over all classes.
- *Experiment 1*. Our research dataset of images of regenerating plants features a relatively small number of classes (stem, callus and shoot) found in a limited number of contexts, unlike highly complex datasets that dominate DNN research. Considering this together with reported successes of random forest in segmenting plant tissues [49, 50], we compared the performance of random forests and PSPNet [29] in segmenting our plant images. PSPNet [29] outperformed random forests by a margin of approximately 30% IoU for each class of interest in the validation dataset (Table3). Furthermore, PSPNet yields a lesser difference between training and validation IoU than was seen with random forests, suggesting that the deep architecture of PSPNet [29] better captures high-level features and avoids overfitting.

**Table 3.** Segmentation IoU score (in %) on the training and validation dataset.
- *Experiment 2*. To gain insight into an appropriate number of training samples for use in our workflow, we explored the relationship between the number of training examples and the quality of segmentation. We trained six different models using 20, 40, 60, 80, 100, and 120 images randomly selected from our training set and tested each model’s performance using the same validation set of 20 images. As shown in Fig.13, the Mean IoU increased significantly as the training sample size increased to 40, 60, and 80. As the sample size increases to 100 and further, additional increases in Mean IoU become markedly diminished but may remain significant and valuable for applications that benefit from high precision. In summary, the deep learning models’ performance relies heavily on the size of the training set, and the inclusion of additional images can almost always be expected to improve the quality of segmentation results. These results demonstrate the importance of a substantial training set for semantic segmentation, underscoring the value in developing efficient and intelligent interactive image segmentation algorithms and annotation tools such as IDEAS.

| Method | Dataset | Background | Stem | Callus | Shoot | mIOU |
| --- | --- | --- | --- | --- | --- | --- |
| RF [50] | train. | 94.62% | 69.83% | 38.75% | 40.82% | 61.00% |
| RF [50] | val. | 93.83% | 57.67% | 20.92% | 39.36% | 52.95% |
| PSPNet [29] | train. | 99%.35 | 89.76% | 73.96% | 74.36% | 84.36% |
| PSPNet [29] | val. | 99.36% | 88.14% | 61.95% | 67.40% | 79.21% |

**Figure 13.**
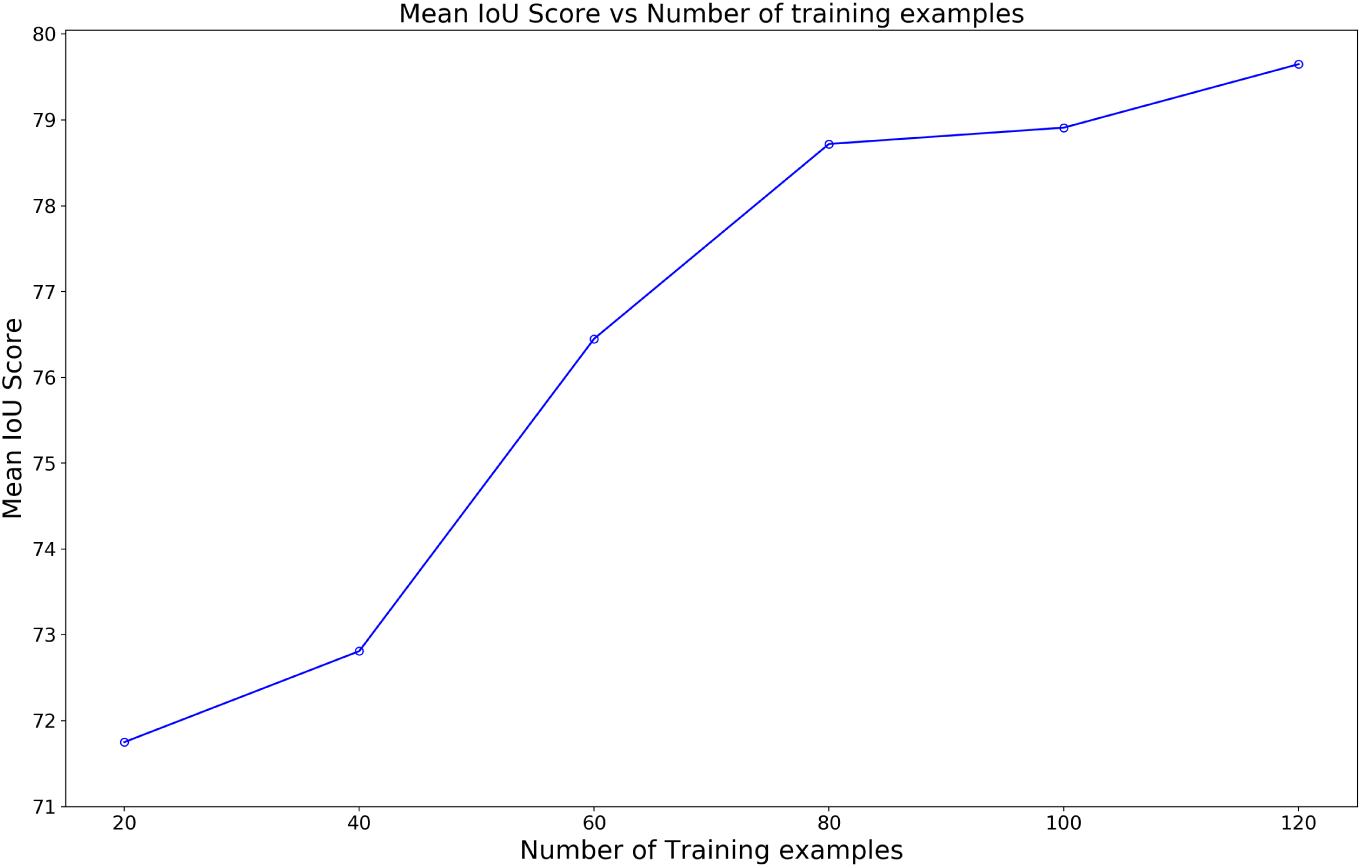
The Mean IoU for different number of training examples.

#### Biological traits

After the semantic segmentation model was trained on the collected training set, we used it to predict labels given unseen images. Next, we calculate the statistic, relative area, which represents the development of specific tissues in each image, and used these in our downstream models for GWAS. Computation of relative area is shown in eq.2:

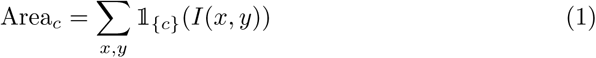

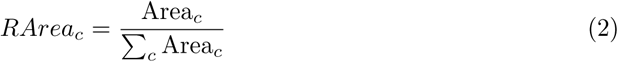

Where *I* is the semantic segmentation from the PSPNet [29] and *c* is the category of the tissue for which relative area is being calculated.

#### Association mapping

We present results from GWAS for a key trait of interest, the area of shoot grown at the final timepoint (fifth week). In this analysis, we apply a mixed linear model approach that is commonly used for GWAS of continuous phenotypes and assumes normality of residuals. To avoid a severe violation of this assumption, genotypes labeled by the model as having zero values (no shoot) were dropped and the remaining phenotype data underwent a natural logarithm transformation. The population analyzed in this case study consists of 326 clones for which genotype data was available and phenotype data is nonzero.

Genotype data used for analysis was obtained from the Bioenergy Science Center at Oak Ridge National Laboratory (https://cbi.ornl.gov/data) and filtered for minor allele frequency and missing rates using PLINK [51]. The filtered dataset used for analysis features approximately 5.3 million single-nucleotide polymorphisms (SNPs), each of which are polymorphic in at least 5% of genotypes and nonmissing in at least 90%.

We employed Genome-wide Efficient Mixed Model Association (GEMMA) [52] as a GWAS method. Mixed models are built for each SNP, explaining the phenotype as a function of the given SNP and covariates. To control for potential confounding factors of cutting size and population stratification, we included covariates of stem diameter, a kinship matrix and three principle components derived from SNPs. Multiple-testing correction performed using the Benjamini-Hochberg method was used to calculate a p-value threshold with a false-discovery rate of 10% [53]. Putative associations with p-values less than or equal to this threshold are reported in Fig.14 and Table.4.

**Figure 14.**
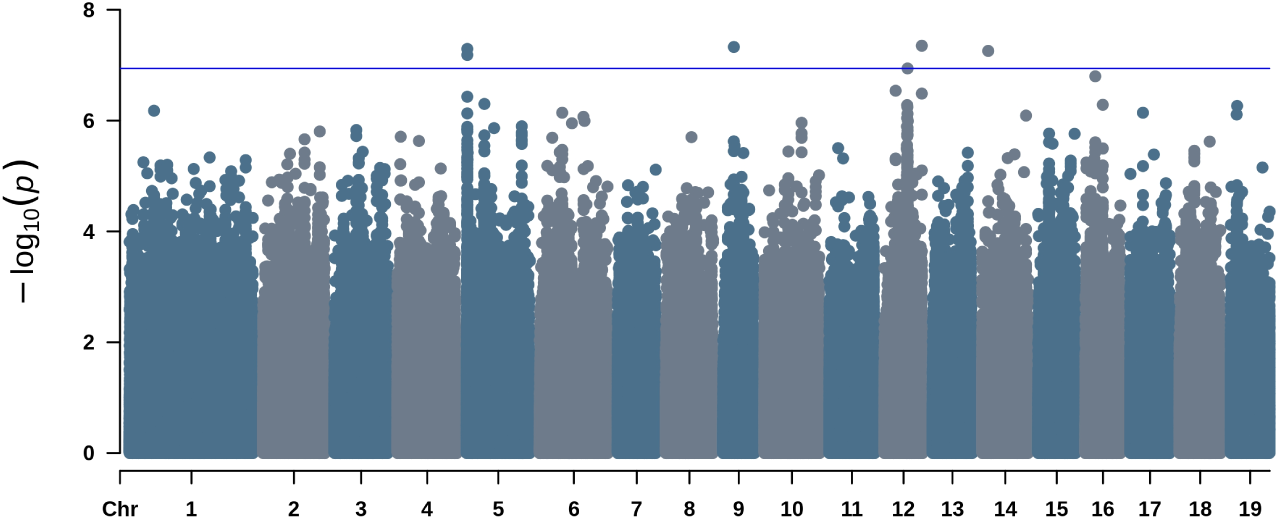
This Manhattan plot shows the significance of genetic markers in models explaining the shoot regeneration phenotype as a function of each marker. The blue line represents a threshold for a false discovery rate of 10%, calculated by the Benjamini-Hochburg method [53].

**Table 4.** QTLs, implicated poplar gene candidates and related Arabidopsis homologs.

| QTL | p-value | Gene candidate | Proximity (bp) | Arabidopsis gene | Similarity |
| --- | --- | --- | --- | --- | --- |
| Chr5:284409 | 5.1e-08 | Potri.005G004700 | 692 | None | N/A |
| Chr5:284964 | 6.5e-08 | Potri.005G004700 | 137 | None | N/A |
| Chr9:4459368 | 4.7e-08 | Potri.009G034700 | 1,283 | AT3G55530: SALT- AND DROUGHT-INDUCED RING FINGER1 | 85.4% |
| Chr12:9509286 | 1.1e-07 | Potri.012G070801 | 8,961 | None | N/A |
| Chr12:15334435 | 4.5e-08 | Potri.012G138800 | 1,155 | AT5G13930: CHALCONE SYNTHASE | 83.0% |
| Chr14:2487581 | 5.5e-08 | Potri.014G029200;<br>Potri.014G029300 | 1,924; 2,556 | AT1G07700: Thioredoxin superfamily protein; AT5G42890: STEROL CAR-RIER PROTEIN 2 | 88.7%;<br>81.3% |

#### Interrogation of putative associations

To determine genes implicated by these genetic markers, we consulted a reference *Populus trichocarpa* genome [54] annotation (version 3.1 available at https://phytozome-next.jgi.doe.gov) to determine relative positions of these genetic markers to gene transcripts. To identify homologs of implicated genes in better-characterized plant species, and thus gain insight into the function of these genes, we referred to a database of Smith-Waterman alignments of predicted peptides between *P. trichocarpa* and the model plant *Arabidopsis thaliana* (https://phytozome-next.jgi.doe.gov).

The two candidate genes implicated by lowest p-values are related to Arabidopsis genes with known roles in regeneration (Fig.14 and Table.4). *CHALCONE SYNTHASE* (*CHS*) encodes a protein essential for a rate-limiting step of the biosynthesis of anthocyanin, which may influence shoot regeneration by dual mechanisms of auxin transport regulation [55] and light stress protection. Loss-of-function mutations of *CHS* are reported to be deficient in shoot regeneration, with a light-dependent effect [56]. *SALT-AND-DROUGHT-INDUCED RING FINGER 1* (*SDIR1*) is an E3 ubiquitin ligase critical for regulation of protein degradation downstream of the hormone abscisic acid. Loss-of-function and overexpression lines of *SDIR1* display enhanced and inhibited levels of *in vitro* seedling germination, respectively [57]. Supplementation of *in vitro* media with abscisic acid has been reported to enhance *in vitro* regeneration in diverse plants, particularly via the route of somatic embryogenesis [58].

Our third candidate (Potri.005G004700) is a gene of unknown function that invites further review by biologists aiming to characterize the genetic basis of plant regeneration. Smith-Waterman alignments reveal no Arabidopsis homolog of this gene.

Table4 shows quantitative trait loci (QTLs) with their p-values calculated by GEMMA. Accession IDs of nearby gene candidates are reported, along with the distance of the QTL from each gene. The most similar Arabidopsis homolog is shown for each candidate gene, along with the similarity as calculated by Smith-Waterman alignment. Chr14:2487581 appears between two genes and both of these possible causative genes are listed. Potri.005G004700 is implicated by two significant QTLs.

## Discussion

While supervised deep learning has demonstrated great power for diverse image tasks, the potential for this technology to be applied to specific tasks is constrained by the abilities of users to produce sufficient training data. Moreover, for plant biology and other fields in which researchers commonly lack a computer science background, an essential prerequisite for the broad dissemination of deep learning is the development of generalizable workflows with user-friendly, high-level interfaces for annotating training data and deploying models. This combination of challenges presents a need for innovation both in algorithm and interface design. To this end,we created a novel annotation system tailored to the needs of plant biologists engaged in phenotyping of plant tissues during any stage of growth or regeneration.

IDEAS enables high-throughput measurement of plant tissues, and can be easily applied to diverse tissue types by user annotation of features of interest. A major advantage of IDEAS is the ability to accurately define tissue object boundaries that are highly complex and thus ignored or misidentified by common annotation methods involving polygons or bounding boxes. Manual pixel-scale annotation can produce highly precise boundaries, but the labor-intensiveness of this process presents an obstacle to generating sizable training sets. Our semantic-guided interactive object segmentation algorithm provides high precision at high speed, delivering substantial reductions to the labor cost of generating high-quality training sets for complex traits.

Worth emphasizing is that interactive segmentation is a more complex problem than semantic segmentation, especially where user annotation involves separate instances of a given type of object (class). Thus, the potential for the semantic prior maps to improve interactive segmentation has several noteworthy limitations. First, in cases where there exist adjacent or touching objects of the same class, the separation of these objects by the user is particularly difficult, and may not be improved by the use of a semantic prior maps lacking instance information. Second, the performance of semantic segmentation tends to be greater for larger objects than for smaller objects. Hence, for small objects, a semantic prior map may provide little or no benefit to the system. These limitations considered, the effective use of a semantic prior map to guide interactive segmentation must include an ability to apply the prior only when it offers a performance benefit. In a preliminary study, we examined the use of prior information and conducted a statistical analysis to determine whether using a semantic prior map helps to accelerate annotation. The semantic prior map was found to confer a statistically significant increase in annotation speed of approximately 20 percent.

Finally, we demonstrated our system using the case study of a GWAS of *in planta* regeneration in poplar. Top associations from this GWAS implicate genes with Arabidopsis homologs that have known roles in regeneration pathways. Regeneration entails hormone-driven processes [59] involving hormones including auxin and abscisic acid, which are regulated by genes including CHS [55] and SDIR1 [57, 58], respectively. The agreement between our results and established models of these genes’ roles in regeneration suggests that our phenotyping system is effective in capturing the shoot regeneration phenotype in our case study. Furthermore, the appearance of an unknown gene as a putative association provides an example of how this system can contribute to genetic discovery.

## Conclusion

We developed a robust deep segmentation phenotyping system utilizing a web-based annotator, IDEAS, for generating ground truth datasets. Using the semantic-guided interactive object segmentation backend, IDEAS provides an accelerated means of labeling objects at pixel-scale with precise boundaries. Using labels generated by the annotator, researchers can train a deep model for semantic segmentation, deploy the model to make predictions over a large dataset and compute statistics summarizing segments of biological interest. Downstream, GWAS revealed genetic markers associated with traits phenotyped by computer vision. Our system can be used by plant biologists who are interested in complex segmentation-based traits, for which generation of large training sets may otherwise be time-prohibitive.

## General

We greatly appreciate the support by Drs. Jerry Tuskan and Wellington Muchero of BESC for their collaboration and advice during research grant development and project execution. We thank the members of the GREAT TREES (Genetic Research on Engineering and Advanced Transformation of Trees) Research Cooperative at Oregon State University for its long term investment in our transformation and regeneration studies. We thank Middleton Spectral Vision (Wisconsin) for their customized imaging system used to capture regeneration data, and high quality imaging system support.

## Poplar SNP dataset

Support for the Poplar GWAS dataset is provided by the U.S. Department of Energy, Office of Science Biological and Environmental Research (BER) via the Bioenergy Science Center (BESC) under Contract No. DE-PS02-06ER64304. The Poplar GWAS Project used resources of the Oak Ridge Leadership Computing Facility and the Compute and Data Environment for Science at Oak Ridge National Laboratory, which is supported by the Office of Science of the U.S. Department of Energy under Contract No. DE-AC05-00OR22725

## Author contributions

JY and DK designed the image analysis pipeline and experimental procedures and performed evaluation of the idea relating to the sections involving computer vision techniques. MN provided important help in the experimental section on GWAS study and writing of this paper. ZZ, NGK, and NAD assisted in the GUI design. EP assisted in collecting data with IDEAS. CM, SS, and FL supervised the project and designed the research.

## Funding

We thank the National Science Foundation Plant Genome Research Program for funding, “Analysis of Genes Affecting Plant Regeneration and Transformation in Poplar.” IOS # 1546900.

## Conflict of interest

There is conflict of interest regarding to this article with Apple Inc., Microsoft Cooperation Inc., and Parametrix.ai Inc..

---

**Figure S1.**
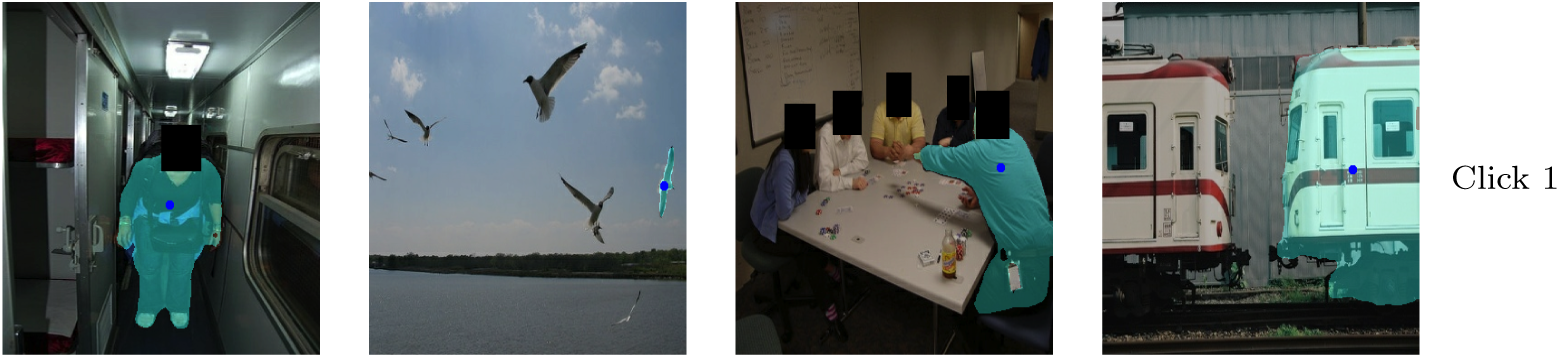
Example predictions on Pascal VOC dataset on a single click.

**Figure S2.**
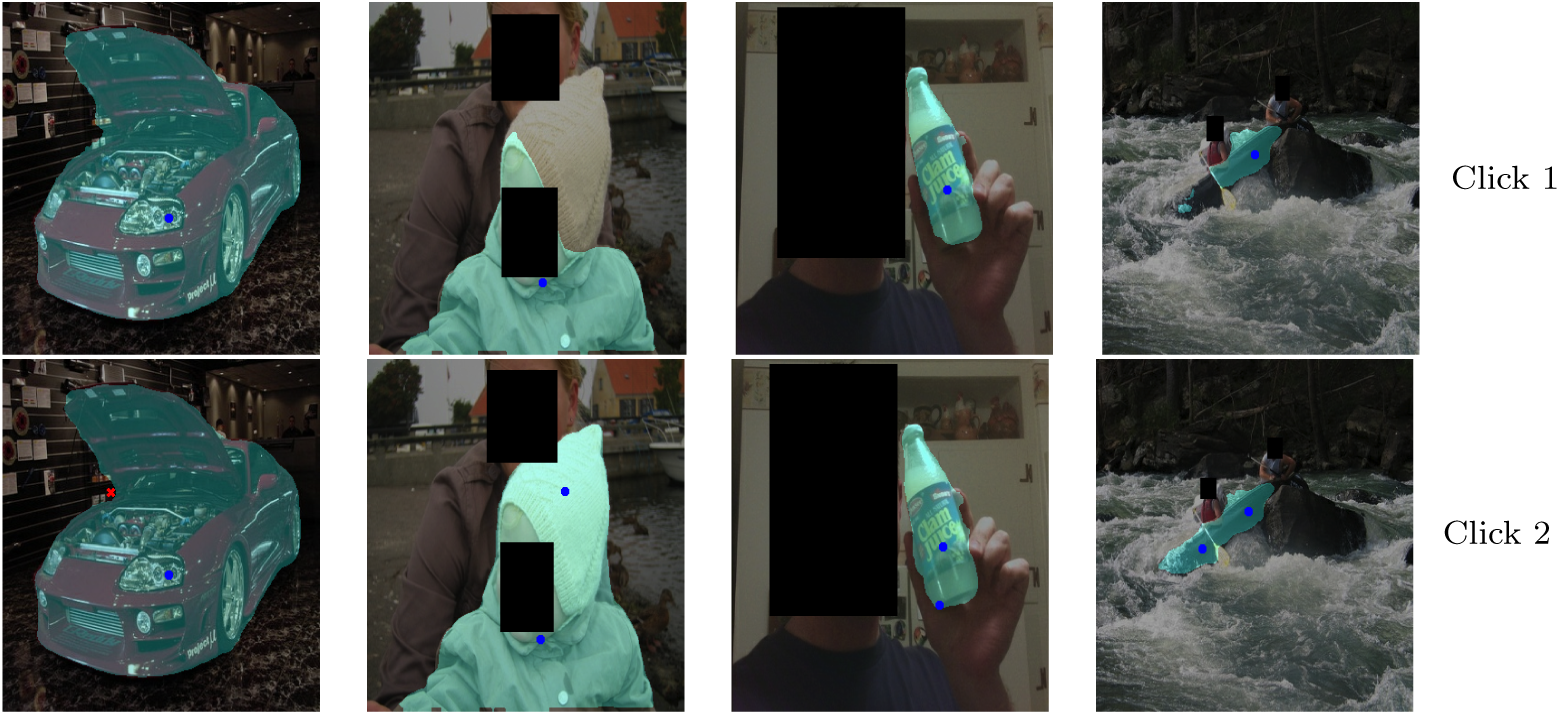
Example predictions on Pascal VOC dataset on two clicks.

**Figure S3.**
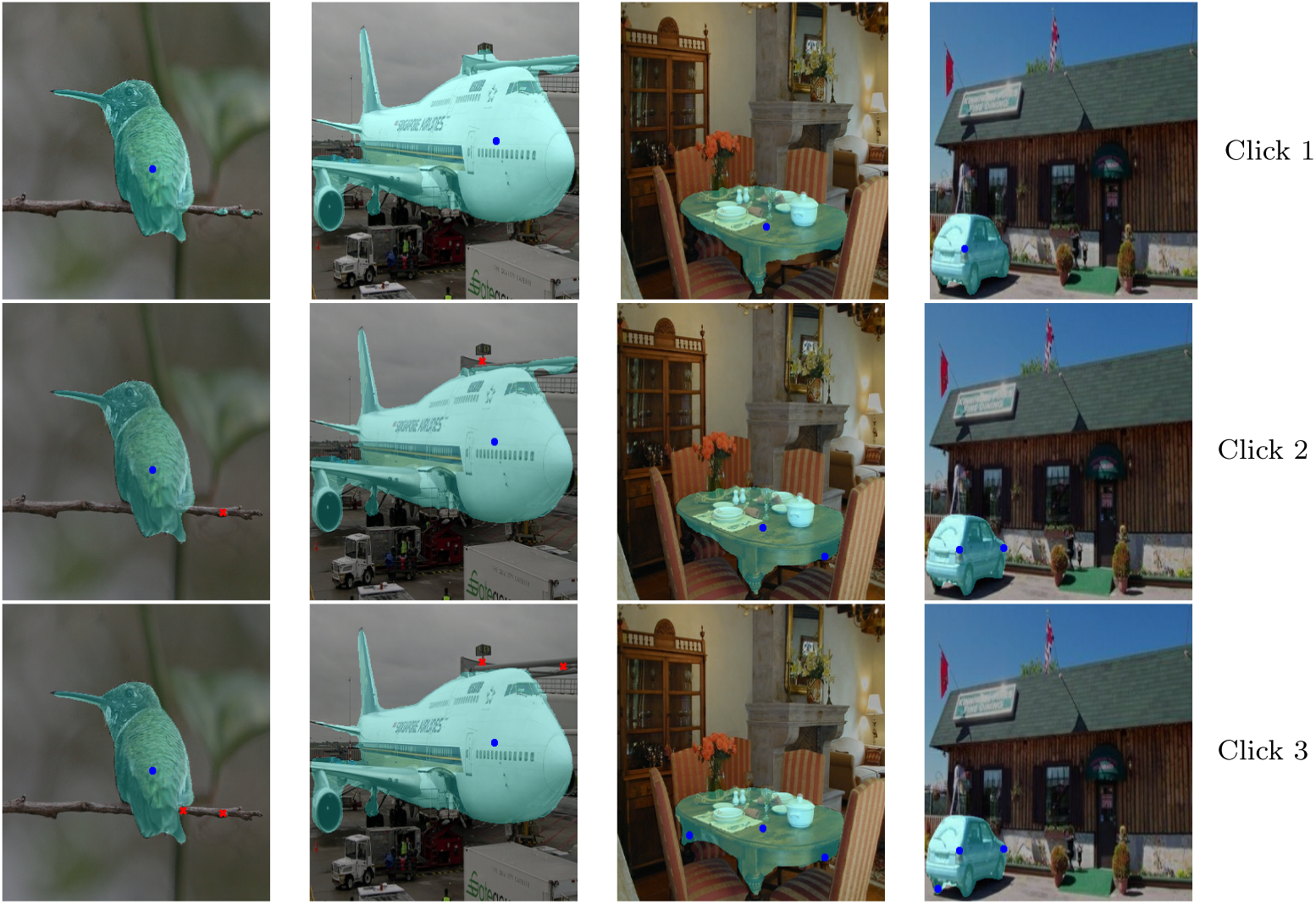
Example predictions on Pascal VOC dataset on three clicks.

**Figure S4.**
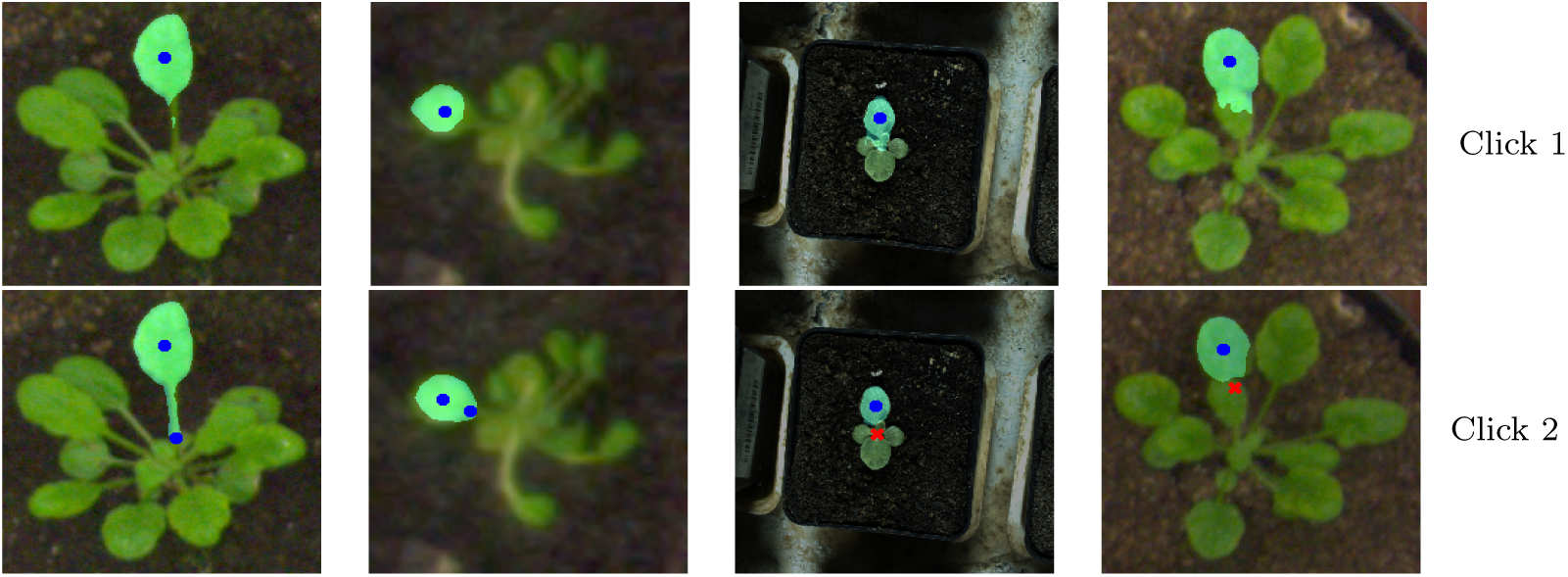
Example predictions on Leaf Segmentation Challenge (LSC) dataset on two clicks.

**Figure S5.**
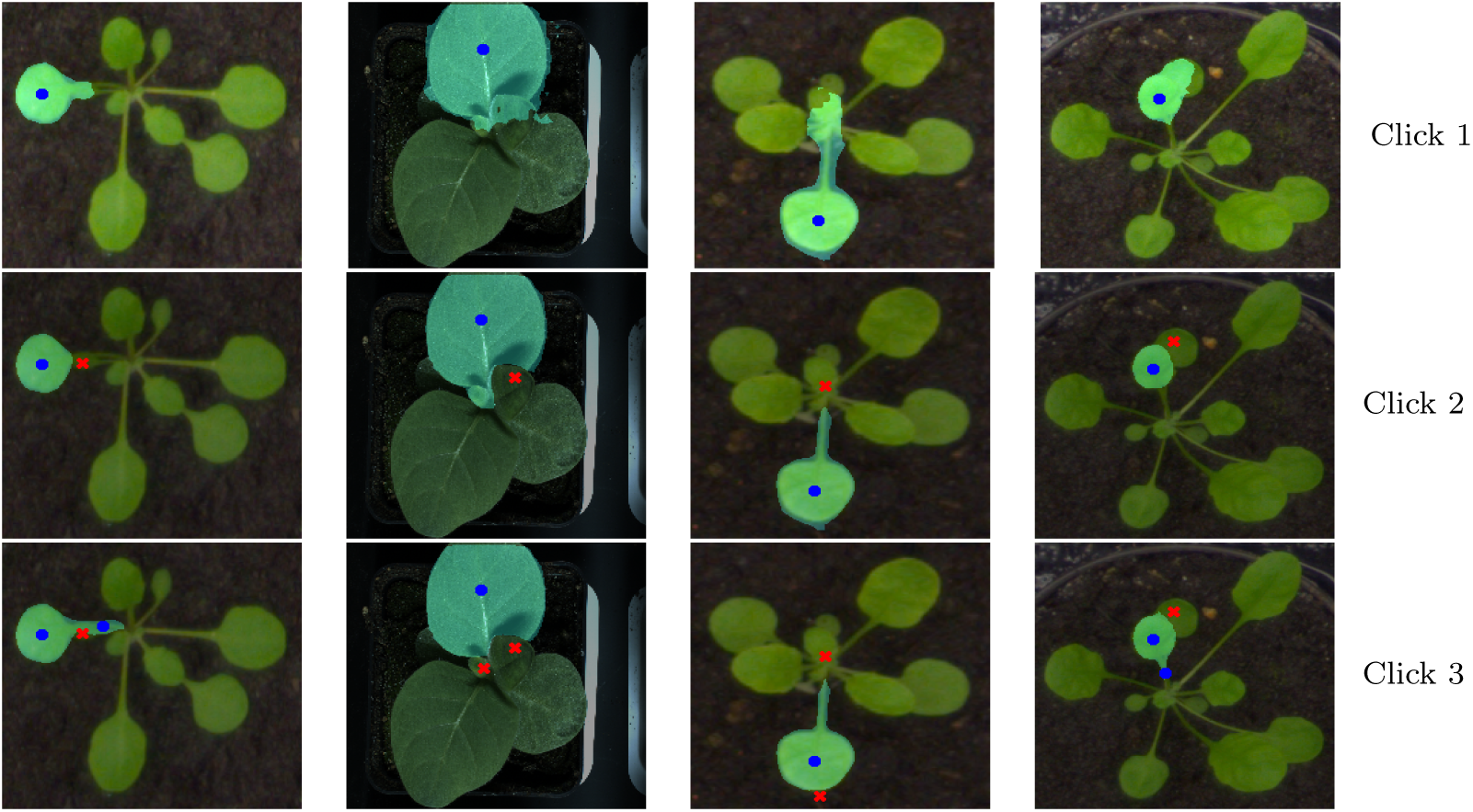
Example predictions on Leaf Segmentation Challenge (LSC) dataset on three clicks.

## Notes

https://ideas.eecs.oregonstate.edu

### Competing Interest Statement

The authors have declared no competing interest.

